# Phylogenetic placement of *Ceratophyllum submersum* based on a complete plastome sequence derived from nanopore long read sequencing data

**DOI:** 10.1101/2023.06.27.546741

**Authors:** Samuel Nestor Meckoni, Benneth Nass, Boas Pucker

## Abstract

**Objective:** Eutrophication poses a mounting concern in today’s world. *Ceratophyllum submersum* L. is one of many plants capable of living in eutrophic conditions, therefore it could play a critical role in addressing the problem of eutrophication. This study aimed to take a first genomic look at *C. submersum*.

**Results:** Sequencing of gDNA from *C. submersum* yielded enough reads to assemble a plastome. Subsequent annotation and phylogenetic analysis validated existing information regarding angiosperm relationships and the positioning of Ceratophylalles in a wider phylogenetic context.

## Introduction

*Ceratophyllum submersum* L., commonly known as soft hornwort, is a subaquatic plant whose genus is the only extant member in the order of Ceratophyllales, placed as a sister clade to the eudicots [1,2]. It is native to Europe, Africa and Asia and grows in stagnant freshwater bodies [3]. Morphological features include long, branching stems that can reach up to several metres in length and leaves that are forked into narrow, filament-like segments that grow in multiple whorls around the stem (Figure 1) [4]. The plant is often green, but can vary in colour from brown to red depending on environmental conditions. Anthocyanins contribute to the colouration of many plant species, and metabolic analyses have detected several derivatives in *C. submersum* [5–7]. It thrives in eutrophic conditions, characterised by low light intensity and high nutrient levels [8,9]. Eutrophication of aquatic environments is indicated by the accumulation of nutrients albeit other parameters and multiple classification systems exist [10]. Due to anthropogenic effects, the occurrences of eutrophic environments are rising which poses a problem e.g. for greenhouse gas emissions [11]. In aquatic systems, eutrophication induces harmful algal blooms (HABs) which are responsible for environmental hazards like the Oder ecological disaster in 2022 [12]. Due to its capabilities, *C. submersum* competes with other phototrophic organisms capable of living in eutrophic conditions. This suggests that it may inhibit the formation of HABs despite its vulnerability to them [7].

**Figure 1:**
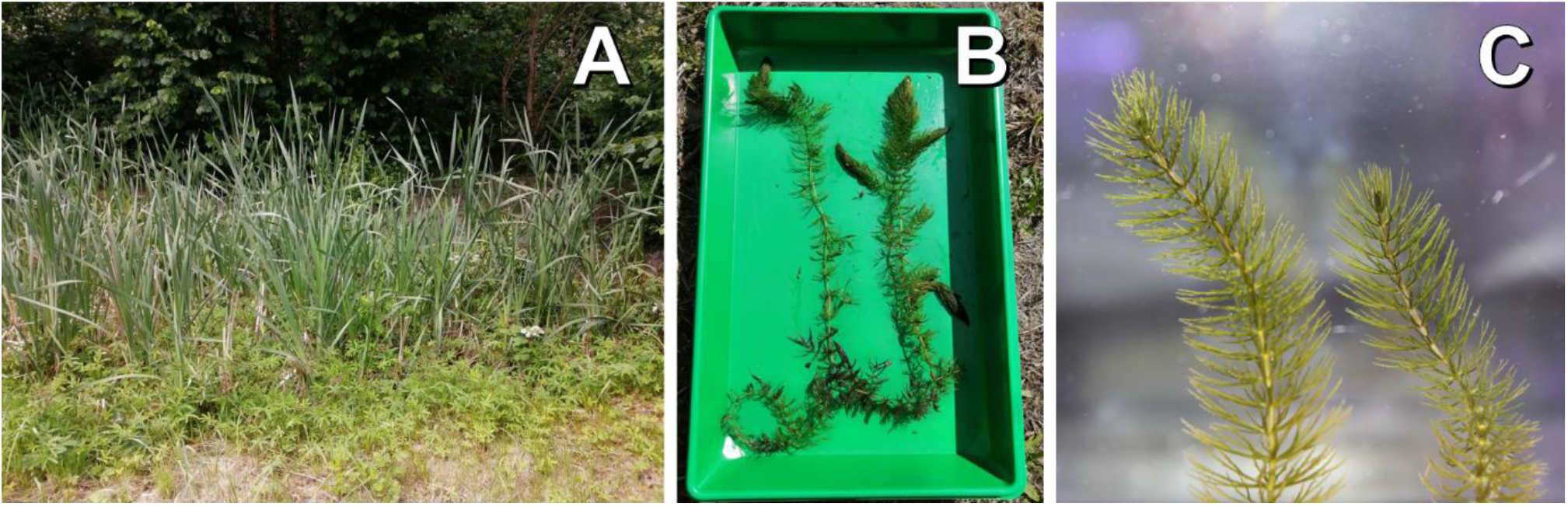
*Ceratophyllum submersum* and its habitat. **A**: The pond (Braunschweig, 52.28062 N / 10.54896 E) where *C. submersum* was collected, **B**: Whole *C. submersum* plant, **C**: Close up view of *C. submersum*.

While a low coverage skimming report has been performed previously [13], the results appear to be missing in the established sequence databases. Here, we utilised nanopore long-read sequencing to obtain the plastome sequence of *C. submersum* and provide a thorough annotation of its genetic content. This analysis allowed us to place the species in a phylogenetic context which informs future studies.

## Main Text

## Results and Discussion

A total of 0.48 Gbp of sequencing data was generated from 94,100 long reads. Out of this, 1,544 reads representing the *C. submersum* plastome were extracted (Additional file 1, PRJEB62706), accounting for 3.3% of the data. Due to the limitations of the Flye assembler used by ptGAUL, only 752 reads (covering about 1.68% of the total data) were utilised for the assembly, as it can only accommodate up to 50x coverage of the estimated assembly size. Considering these factors, only 0.27 Gbp of sequencing data are required to assemble an approximately 160 kbp sized plastome, provided that the ratio of plastid DNA to nuclear DNA does not exceed 3%. We calculated that about 0.8 Gbp of total genomic sequencing data would be adequate for cases where the plastid DNA content is even lower (1%). High-quality assemblies can be achieved with coverage levels lower than 50, so a smaller amount of data may still be sufficient. We inferred a sequencing goal of 1 Gbp to enable the assembly of a plastome sequence without prior plastid purification. If fresh leaf material is chosen, a higher plastid DNA portion should be achievable, potentially reducing the amount of data needed for successful assembly.

The plastome assembly of *C. submersum* resulted in a 155,767 bp long sequence with a GC content of 38.26%. In total, 75 unique protein coding genes were annotated (Additional file 2). The *C. submersum* plastome sequence is 485 bp shorter than the reference plastome sequence of *Ceratophyllum demersum* L. that has a length of 156,252 bp [14]. While the GC content is similar (*C. demersum*: 38.22, *C. submersum*: 38.26) the amount of unique annotated genes in *C. submersum* is smaller than in *C. demersum* (75 against 79). After the annotation some predicted coding sequences were flawed (e.g. containing multiple stop codons). Manual evaluation revealed four homopolymeric regions in which a frameshift would correct the annotation. Since homopolymers are a frequent error type in ONT reads, manual correction in those regions is appropriate [15]. The comparison of those regions to all plastome reads suggested a correction of two of these regions, namely the adenine at position 3,094 and the thymines at position 3,143, at position 86,261, and at position 86,262 were inserted.

The sequencing took place on a flow cell that was previously used to analyse gDNA from *Digitalis purpurea* L. Therefore, a phylogenetic tree was calculated to validate clean and distinguishable sequencing data, incorporating our plastome assemblies for both species (*D. purpurea* data: PRJEB62706) (Figure 2).

**Figure 2:**
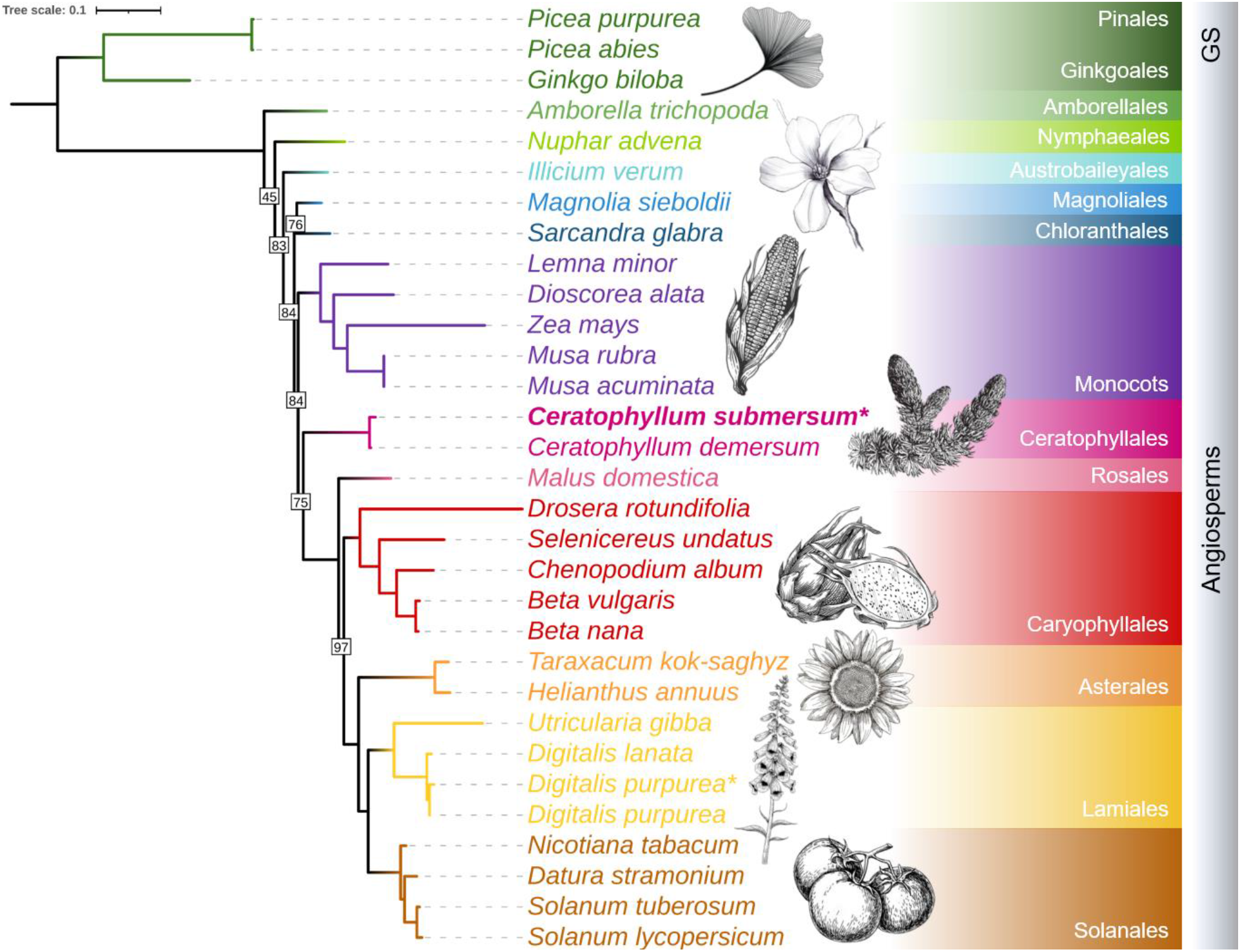
Phylogenetic tree of selected spermatophytes based on plastome protein sequences. The asterisks (*) indicate plastome assemblies generated in this study. Within the angiosperms, the orders are differently coloured. The eudicotyledons are the only exception as they are represented by several orders. All nodes received full bootstrapping support (100%), except those displaying the actual value. Please see the material and methods section for further description of the tree calculation. Visualisation was done in iTOL 6.7.6 [16]. The full tree including the outgroup *Chlamydomonas reinhardtii* can be found in the additional files (Additional file 4). GS = gymnosperms. The sketches are licensed by adobe and depositphotos (Standard License: https://wwwimages2.adobe.com/content/dam/cc/en/legal/servicetou/Stock-Additional-Terms_en_US_20221205.pdf, https://depositphotos.com/license.html).

Multiple plastome reference sequences were chosen to represent the angiosperm clade as well as *Chlamydomonas reinhardtii* as outgroup (for full list and references see Additional file 3). The phylogenetic tree (figure 2, for full tree see Additional file 4) classifies *C. submersum* close to its reference *C. demersum* and our *D. purpurea* plastome assembly close to the RefSeq *D. purpurea* plastome generated by Zhao *et al*., 2023 [17]. The angiosperm clade is represented in accordance with the current APG IV classification except for the exact separation/placement of the Magnoliids and the Chloranthales, which is still controversial [2]. This underlines the significance of plastome sequences for modern plant phylogenetics [18]. The plastome assembly and phylogenetic analysis presented in this study provides first steps towards genetic and genomic characterization of *Ceratophyllum submersum*. Further research is needed to determine its nuclear genome and to explore potential applications of this plant, such as its use as a valuable resource or as an agent to mitigate environmental hazards.

## Material and Methods

### Plant Material, gDNA extraction and sequencing

*Ceratophyllum submersum* was collected from a small pond in Braunschweig (52.28062 N / 10.54896 E) and kept in cultivation at the Institute of Plant Biology at TU Braunschweig. The artificial pond needs occasional water replenishment. Foliage from surrounding shrubs and dead aquatic plants lead to a high eutrophy (see figure 1A). *C. submersum* shoot tips (figure 1C) were harvested for gDNA extraction conducted with a CTAB method [19–21]. Short DNA fragments were depleted with the Short Read Eliminator kit (Pacific Biosciences). Library preparation for ONT sequencing was started with 1μg of DNA that was first repaired with the NEBNext^®^ Companion Module and then processed according to the SQK-LSK109 protocol (Oxford Nanopore Technologies). For ONT sequencing, a R9.4.1 flow cell was used with a MinION. ONT sequencing is one of the leading sequencing technologies in plant genomics [22], and was applied in this project to generate a complete plastome sequence based on long reads. Prior to *C. submersum* sequencing, the flow cell was already utilised for *D. purpurea* gDNA sequencing. Processing of raw data was performed with guppy v6.4.6+ae70e8f (https://community.nanoporetech.com) which internally called minimap2 v2.24-r1122 [23] with default parameters on a graphical processor unit (GPU) in the de.NBI cloud to generate FASTQ files. Guppy was run with the default configuration of dna_r9.4.1_450bps_hac.

### Plastome Assembly and Annotation

The FASTQ files were subjected to a plastome assembly with ptGAUL using standard parameters [23–26]. As reference the *C. demersum* plastome from the NCBI RefSeq database (release 216) was used [14,27]. Since the ptGAUL results consist of two assemblies (one per path) the OGDRAW plastome maps were compared between the reference and the assemblies to decide which one is closer to the reference. The *C. submersum* assembly needed to be reverse complemented and the sequence start was adjusted to the *C. demersum* sequence. GC contents of the assembly were calculated using the contig_stats.py script [28]. For annotation, GeSeq was used (see Additional file 5 for parameters) [29–38]. Protein coding genes were extracted from the resulting GBSON.json file using our own script JSON_2_peptide_fasta_v1.0.py [39]. Frameshifts in the coding sequences of incorrectly translated peptide sequences were identified in two steps. First, all open reading frames (ORFs) were annotated by EMBOSS sixpack. Then, NCBI blastx results were mapped against the ORFs to specify the location of the possible frameshifts [40,41]. Further evaluation of these frameshifts was conducted with the help of a read mapping. All plastome reads (FASTQ) were mapped against the plastome sequence (FASTA) with minimap2 (v2.24-r1122, parameters: -ax map-ont --secondary=no -t 27) to generate a SAM file [23]. From that, a BAM file and its corresponding index file was generated with samtools 1.10 [42]. The mapping was then analysed in the Integrative Genomics Viewer (IGV) 2.16.1 [43]. The assembly underwent correction only if supported by more than a quarter and a minimum of ten reads.

Data preparation prior to submission was performed by extracting all the read IDs from the ‘new_filter_gt3000.fa’, generated by ptGAUL, which contains all the identified plastome reads. The plastome reads were extracted from the FAST5 and FASTQ raw read datasets via the ‘fast5_subset’ command from the ont_fast5_api tool and our custom script FASTQ_extractor_from_FAST5_mapping_file.py (both using default parameters) [39,44].

### Phylogenetic analysis

Reference data retrieval and phylogenetic supermatrix tree construction was performed by our newly developed Python pipeline PAPAplastomes (Pipeline for the Automatic Phylogenetic Analysis of plastomes). It integrates established external tools for complex steps (Additional file 6) [45]. Reference species, closely related species of interest, and outgroup species can be specified via a config file. First, NCBI RefSeq plastome data is downloaded and the reference plastome peptide sequences are extracted from this collection. These peptide sequences are further combined with the peptide sequences derived from the assemblies representing plastomes of interest. The pre-OrthoFinder trimming step removes potential paralogs with the exact same sequence, sequences which contain asterisks, and sequences that are shorter than 10 amino acids (this threshold value is adjustable). Next, OrthoFinder v2.5.4 [46–49] is applied. Post-Orthofinder processing includes four steps. Removal of outlier sequences (first step), deletion of orthogroups missing the species of interest or their references (second step), removal of orthogroups harbouring fewer species than the pre-defined outgroup species (third step), and paralog cleaning (fourth step). These steps are explained in more detail below. **First step:** Since the OrthoFinder results include phylogenetic trees of each orthogroup, outlier identification is conducted with the help of the Python module Dendropy v4.5.2 [50]. Per orthogroup the edge length for each taxon is accessed except for outgroup species. Based on these lengths, outliers are identified by the 1.5*IQR method (inter quartile range) i.e. sequences with a distance larger than 1.5 times the variation are excluded. **Second step:** The species of an orthogroup are listed and if pre-defined species are all either present or absent, the orthogroup will be kept. Otherwise, the orthogroup will be discarded. This is suggested for closely related species i.e. from the same genus and can be specified by the user in the config file. **Third step:** Orthogroups consisting solely of the outgroup species are exempt from this criterion. **Fourth step:** Among remaining paralogs within one species, only the longest sequence is kept.

The cleaned orthogroups are then aligned with MAFFT v7.453 (--maxiterate 1000 --localpair) [51,52] and alignments are concatenated. Phylogenetic supermatrix tree calculation is performed by IQ-TREE (multicore version 1.6.12 for Linux 64-bit built Aug 15 2019; with the ‘-nt AUTO -bb 1000’ options, seed: 291752) based on the concatenated alignment of all orthogroups [53–55].

## Limitations

Before sequencing *C. submersum*, the flow cell had already been utilised for sequencing gDNA of *D. purpurea*. Insufficient DNA availability hindered the complete sequencing and assembly of *C. submersum*’s nuclear genome. Future optimisations in DNA extraction methods dedicated to small aquatic plants could overcome this limitation.

## Abbreviations

CDS: coding sequence
gDNA: genomic DNA
GPU: graphical processing unit
GS: gymnosperm
HAB: harmful algal bloom
RefSeq: NCBI Reference Sequence Database
IQR: interquartile range
ONT: Oxford Nanopore Technologies

## Declarations

### Ethics approval and consent to participate

Not applicable.

### Consent for publication

Not applicable.

### Availability of data and material

The datasets generated and analysed during the current study are available under PRJEB62706. Scripts developed for the data analysis are available from Codeberg (https://codeberg.org/snmeckoni/scripts, https://codeberg.org/snmeckoni/PAPAplastomes) and GitHub (https://github.com/bpucker/GenomeAssembly).

### Competing interests

The authors declare that they have no conflict of interest.

### Funding

Not applicable.

### Author’s contributions

Samuel Nestor Meckoni, Benneth Nass and Boas Pucker contributed to the concept and design of the studies, wet and dry lab work and manuscript writing. The authors have read the final version of the manuscript and agree to its submission to BMC Research Notes.

## Acknowledgments

We thank Dr. Christiane Evers (Institute for Plant Biology, TU Braunschweig) and Walter Wimmer (Lower Saxony Department for Water, Coastal and Nature Conservation) for excellent support in taxonomically classifying several specimens. We are also grateful to Sarah Winnier for improving the language of this article. This work was supported by the BMBF-funded de.NBI Cloud within the German Network for Bioinformatics Infrastructure (031A532B, 031A533A, 031A533B, 031A534A, 031A535A, 031A537A, 031A537B, 031A537C, 031A537D, 031A538A). We acknowledge support by the Open Access Publication Funds of Technische Universität Braunschweig.

## Legends

**Additional file 1: FASTQ stats of the *Ceratophyllum submersum* plastome reads**. The stats were calculated with the help of the script FASTQ_stats3.py (https://github.com/bpucker/GenomeAssembly). (Additional file 1.txt)

**Additional file 2: *Ceratophyllum submersum* plastome map derived from OGDRAW**. (Additional file 2.pdf)

**Additional file 3: List of references of the reference plastomes**. The data was extracted by PAPAplastomes and originates from the RefSeq database. (Additional file 3.txt)

**Additional file 4: Full phylogenetic tree with outgroup species *Chlamydomonas reinhardtii***. (Additional file 4.pdf)

**Additional file 5: Exact parameters for the GeSeq run**. Some parameters from the online tool GeSeq consist of a checkbox that has to be checked. Therefore, the value “check” is stated. (Additional file 5.xlsx)

**Additional file 6: PAPAplastome workflow chart**. (Additional file 6.pdf)

## Notes

https://codeberg.org/snmeckoni/PAPAplastomes

https://codeberg.org/snmeckoni/scripts

https://github.com/bpucker/GenomeAssembly

### Competing Interest Statement

The authors have declared no competing interest.

### Summary of Updates

We present a revised version of our manuscript, "Phylogenetic placement of Ceratophyllum submersum based on a complete plastome sequence derived from nanopore long read sequencing data", reflecting extensive revisions resulting from the peer review process. We appreciate the valuable feedback from the reviewers, these changes improve the quality and clarity of our work.

## References

1. Les DH. The Origin and Affinities of the Ceratophyllaceae. Taxon. 1988 May;37(2):326–45.

2. The Angiosperm Phylogeny Group, Chase MW, Christenhusz MJM, Fay MF, Byng JW, Judd WS, et al. An update of the Angiosperm Phylogeny Group classification for the orders and families of flowering plants: APG IV. Bot J Linn Soc. 2016 May 1;181(1):1–20.

3. POWO. Plants of the World Online. Facilitated by the Royal Botanic Gardens, Kew. [Internet]. Plants of the World Online. 2023 [cited 2023 Apr 16]. Available from: http://powo.science.kew.org/taxon/urn:lsid:ipni.org:names:163088-1

4. Düll R, Kutzelnigg H. Taschenlexikon der Pflanzen Deutschlands und angrenzender Länder: die häufigsten mitteleuropäischen Arten im Porträt. 7th ed. Wiebelsheim: Quelle & Meyer; 2011. 932 p. (Quelle & Meyer Taschenlexikon).

5. Les DH. Systematics and Evolution of Ceratophyllum L. (Ceratophyllaceae): A Monograph (Taxonomy, Flavonoids, Aquatic plants, Phytogeography, Variation). [Columbus]: The Ohio State University; 1986.

6. Les DH. Ceratophyllaceae. In: Kubitzki K, Rohwer JG, Bittrich V, editors. Flowering Plants · Dicotyledons [Internet]. Berlin, Heidelberg: Springer Berlin Heidelberg; 1993 [cited 2023 Apr 16]. p. 246–50. Available from: http://link.springer.com/10.1007/978-3-662-02899-5_24

7. Ujvárosi AZ, Riba M, Garda T, Gyémánt G, Vereb G, M-Hamvas M, et al. Attack of Microcystis aeruginosa bloom on a Ceratophyllum submersum field: Ecotoxicological measurements in real environment with real microcystin exposure. Sci Total Environ. 2019 Apr; 662:735–45.

8. Zartes Hornblatt (Ceratophyllum submersum) [Internet]. [cited 2023 Apr 5]. Available from: https://www.oekologie-seite.de/index.php?id=24&pid=642

9. Ellenberg H. Vegetation Mitteleuropas mit den Alpen. 6., vollständig neu bearbeitete und stark erweiterte Auflage. Leuschner C, Dierschke H, editors. Verlag Eugen Ulmer; 2010.

10. Ayele HS, Atlabachew M. Review of characterization, factors, impacts, and solutions of Lake eutrophication: lesson for lake Tana, Ethiopia. Environ Sci Pollut Res. 2021 Mar;28(12):14233–52.

11. Li Y, Shang J, Zhang C, Zhang W, Niu L, Wang L, et al. The role of freshwater eutrophication in greenhouse gas emissions: A review. Sci Total Environ. 2021 May; 768:144582.

12. European Commission: Joint Research Centre. An EU analysis of the ecological disaster in the Oder River of 2022: lessons learned and research based recommendations to avoid future ecological damage in EU rivers, a joint analysis from DG ENV, JRC and the EEA. [Internet]. Publications Office of the European Union; 2023 [cited 2023 Apr 16]. Available from: https://data.europa.eu/doi/10.2760/067386

13. Alsos IG, Lavergne S, Merkel MKF, Boleda M, Lammers Y, Alberti A, et al. The Treasure Vault Can be Opened: Large-Scale Genome Skimming Works Well Using Herbarium and Silica Gel Dried Material. Plants. 2020 Apr 1;9(4):432.

14. Moore MJ, Bell CD, Soltis PS, Soltis DE. Using plastid genome-scale data to resolve enigmatic relationships among basal angiosperms. Proc Natl Acad Sci. 2007 Dec 4;104(49):19363–8.

15. Delahaye C, Nicolas J. Sequencing DNA with nanopores: Troubles and biases. PLOS ONE. 2021 Oct 1;16(10):e0257521.

16. Letunic I, Bork P. Interactive Tree Of Life (iTOL) v5: an online tool for phylogenetic tree display and annotation. Nucleic Acids Res. 2021 Jul 2;49(W1):W293–6.

17. Zhao F, Liu B, Liu S, Min DZ, Zhang T, Cai J, et al. Disentangling a 40-year-old taxonomic puzzle: the phylogenetic position of Mimulicalyx (Lamiales). Bot J Linn Soc. 2023 Jan 25;201(2):135–53.

18. Gitzendanner MA, Soltis PS, Yi TS, Li DZ, Soltis DE. Plastome Phylogenetics: 30 Years of Inferences Into Plant Evolution. In: Advances in Botanical Research [Internet]. Elsevier; 2018 [cited 2022 Aug 9]. p. 293–313. Available from: https://linkinghub.elsevier.com/retrieve/pii/S0065229617300885

19. Pucker B. Plant DNA extraction and preparation for ONT sequencing. 2020 Mar 21 [cited 2023 Feb 14]; Available from: https://dx.doi.org/10.17504/protocols.io.bcvyiw7w

20. Siadjeu C, Pucker B, Viehöver P, Albach DC, Weisshaar B. High Contiguity de novo Genome Sequence Assembly of Trifoliate Yam (Dioscorea dumetorum) Using Long Read Sequencing. Genes. 2020 Mar;11(3):274.

21. Pucker B, Rückert C, Stracke R, Viehöver P, Kalinowski J, Weisshaar B. Twenty-Five Years of Propagation in Suspension Cell Culture Results in Substantial Alterations of the Arabidopsis Thaliana Genome. Genes. 2019 Sep;10(9):671.

22. Pucker B, Irisarri I, de Vries J, Xu B. Plant genome sequence assembly in the era of long reads: Progress, challenges and future directions. Quant Plant Biol. 2022;3:e5.

23. Li H. Minimap2: pairwise alignment for nucleotide sequences. Birol I, editor. Bioinformatics. 2018 Sep 15;34(18):3094–100.

24. Zhou W, Armijos C, Lee C, Lu R, Wang J, Ruhlman T, et al. Plastid Genome Assembly Using Long-read Data (ptGAUL) [Internet]. bioRxiv; 2022 [cited 2022 Nov 23]. p. 2022.11.19.517194. Available from: https://www.biorxiv.org/content/10.1101/2022.11.19.517194v1

25. Shen W, Le S, Li Y, Hu F. SeqKit: A Cross-Platform and Ultrafast Toolkit for FASTA/Q File Manipulation. Zou Q, editor. PLOS ONE. 2016 Oct 5;11(10):e0163962.

26. Kolmogorov M, Yuan J, Lin Y, Pevzner PA. Assembly of long, error-prone reads using repeat graphs. Nat Biotechnol. 2019 May;37(5):540–6.

27. O’Leary NA, Wright MW, Brister JR, Ciufo S, Haddad D, McVeigh R, et al. Reference sequence (RefSeq) database at NCBI: current status, taxonomic expansion, and functional annotation. Nucleic Acids Res. 2016 Jan 4;44(D1):D733–45.

28. Pucker B, Holtgräwe D, Rosleff Sörensen T, Stracke R, Viehöver P, Weisshaar B. A De Novo Genome Sequence Assembly of the Arabidopsis thaliana Accession Niederzenz-1 Displays Presence/Absence Variation and Strong Synteny. Vandepoele K, editor. PLOS ONE. 2016 Oct 6;11(10):e0164321.

29. Tillich M, Lehwark P, Pellizzer T, Ulbricht-Jones ES, Fischer A, Bock R, et al. GeSeq – versatile and accurate annotation of organelle genomes. Nucleic Acids Res. 2017 Jul 3;45(W1):W6–11.

30. Laslett D. ARAGORN, a program to detect tRNA genes and tmRNA genes in nucleotide sequences. Nucleic Acids Res. 2004 Jan 2;32(1):11–6.

31. Kent WJ. BLAT —The BLAST -Like Alignment Tool. Genome Res. 2002 Apr 1;12(4):656–64.

32. Ian Small, Ian Castleden. Chloë: Organelle Annotator [Internet]. 2020. Available from: https://github.com/ian-small/chloe

33. Nawrocki EP, Eddy SR. Infernal 1.1: 100-fold faster RNA homology searches. Bioinformatics. 2013 Nov 15;29(22):2933–5.

34. Wheeler TJ, Eddy SR. nhmmer: DNA homology search with profile HMMs. Bioinformatics. 2013 Oct 1;29(19):2487–9.

35. Edgar RC. MUSCLE: multiple sequence alignment with high accuracy and high throughput. Nucleic Acids Res. 2004 Mar 1;32(5):1792–7.

36. Greiner S, Lehwark P, Bock R. OrganellarGenomeDRAW (OGDRAW) version 1.3.1: expanded toolkit for the graphical visualization of organellar genomes. Nucleic Acids Res. 2019 Jul 2;47(W1):W59–64.

37. Abascal F, Zardoya R, Telford MJ. TranslatorX: multiple alignment of nucleotide sequences guided by amino acid translations. Nucleic Acids Res. 2010 Jul 1;38(suppl_2):W7–13.

38. Chan PP, Lin BY, Mak AJ, Lowe TM. tRNAscan-SE 2.0: improved detection and functional classification of transfer RNA genes. Nucleic Acids Res. 2021 Sep 20;49(16):9077–96.

39. Meckoni SN. small scripts codeberg repository [Internet]. 2023 [cited 2023 May 30]. Available from: https://codeberg.org/snmeckoni/scripts

40. Madeira F, Pearce M, Tivey ARN, Basutkar P, Lee J, Edbali O, et al. Search and sequence analysis tools services from EMBL-EBI in 2022. Nucleic Acids Res. 2022 Jul 1;50(W1):W276–9.

41. NCBI blastx [Internet]. Bethesda (MD): National Library of Medicine (US), National Center for Biotechnology Information; [cited 2023 Jun 8]. Available from: https://blast.ncbi.nlm.nih.gov/Blast.cgi?PROGRAM=blastx&PAGE_TYPE=BlastSearch&LINK_LOC=blasthome

42. Li H, Handsaker B, Wysoker A, Fennell T, Ruan J, Homer N, et al. The Sequence Alignment/Map format and SAMtools. Bioinformatics. 2009 Aug 15;25(16):2078–9.

43. Robinson JT, Thorvaldsdóttir H, Winckler W, Guttman M, Lander ES, Getz G, et al. Integrative genomics viewer. Nat Biotechnol. 2011 Jan;29(1):24–6.

44. ont_fast5_api github repository [Internet]. nanoporetech; 2022 [cited 2023 May 30]. Available from: https://github.com/nanoporetech/ont_fast5_api

45. Meckoni SN. PAPAplastomes [Internet]. 2023 [cited 2023 Jun 26]. Available from: https://codeberg.org/snmeckoni/PAPAplastomes

46. Emms DM, Kelly S. OrthoFinder: phylogenetic orthology inference for comparative genomics. Genome Biol. 2019 Dec;20(1):238.

47. Emms DM, Kelly S. OrthoFinder: solving fundamental biases in whole genome comparisons dramatically improves orthogroup inference accuracy. Genome Biol. 2015 Dec;16(1):157.

48. Emms DM, Kelly S. STRIDE: Species Tree Root Inference from Gene Duplication Events. Mol Biol Evol. 2017 Dec 1;34(12):3267–78.

49. Emms DM, Kelly S. STAG: Species Tree Inference from All Genes [Internet]. Evolutionary Biology; 2018 Feb [cited 2023 Mar 20]. Available from: http://biorxiv.org/lookup/doi/10.1101/267914

50. Sukumaran J, Holder MT. DendroPy: a Python library for phylogenetic computing. Bioinformatics. 2010 Jun 15;26(12):1569–71.

51. Katoh K. MAFFT: a novel method for rapid multiple sequence alignment based on fast Fourier transform. Nucleic Acids Res. 2002 Jul 15;30(14):3059–66.

52. Katoh K, Standley DM. MAFFT Multiple Sequence Alignment Software Version 7: Improvements in Performance and Usability. Mol Biol Evol. 2013 Apr 1;30(4):772–80.

53. Nguyen LT, Schmidt HA, von Haeseler A, Minh BQ. IQ-TREE: A Fast and Effective Stochastic Algorithm for Estimating Maximum-Likelihood Phylogenies. Mol Biol Evol. 2015 Jan;32(1):268–74.

54. Kalyaanamoorthy S, Minh BQ, Wong TKF, von Haeseler A, Jermiin LS. ModelFinder: fast model selection for accurate phylogenetic estimates. Nat Methods. 2017 Jun;14(6):587–9.

55. Minh BQ, Nguyen MAT, von Haeseler A. Ultrafast Approximation for Phylogenetic Bootstrap. Mol Biol Evol. 2013 May 1;30(5):1188–95.

