## Supplementary figures and images for "Phylogenetic placement of *Ceratophyllum submersum* based on a complete plastome sequence derived from nanopore long read sequencing data"

### Additional file 2

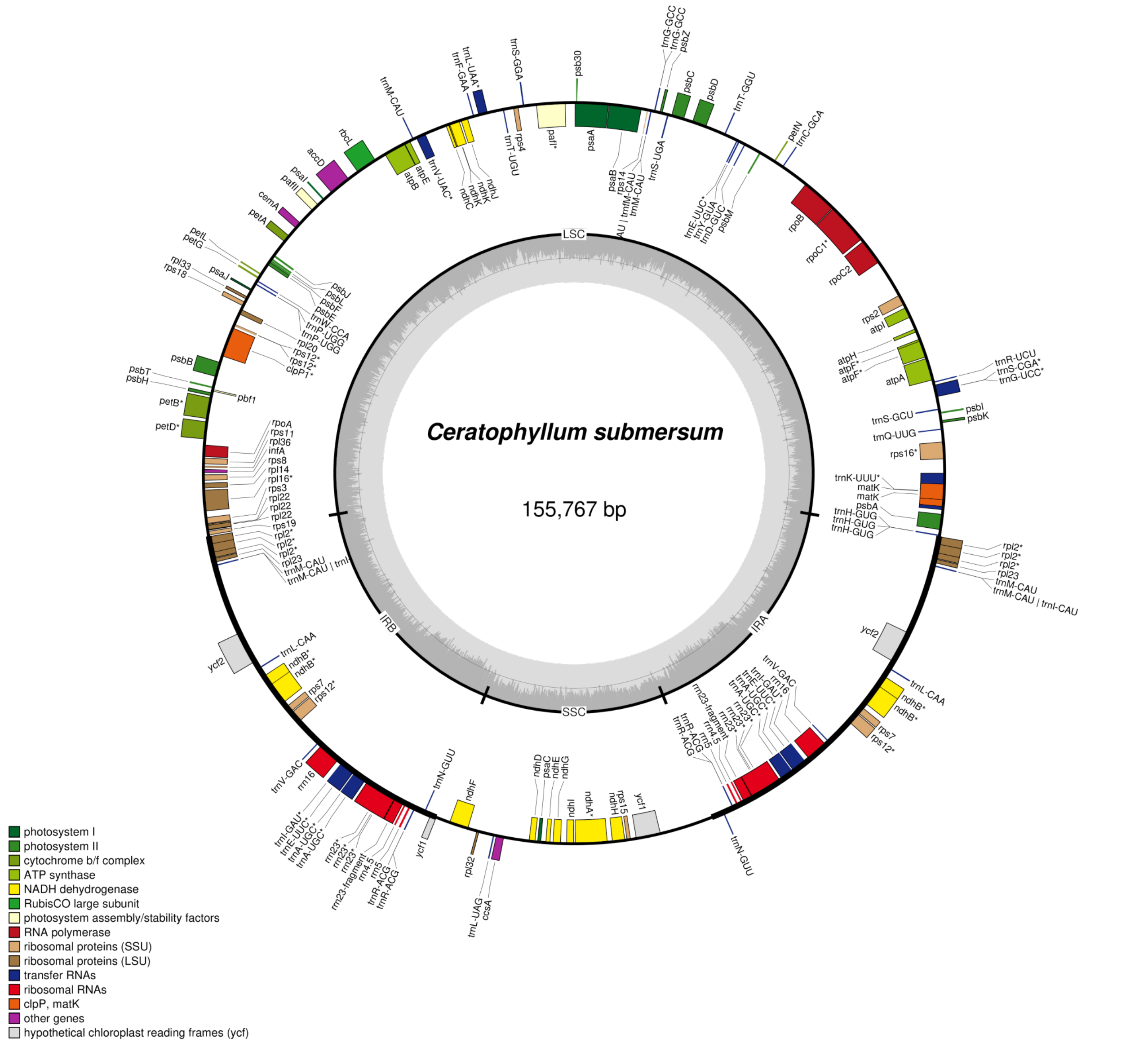

### Additional file 4

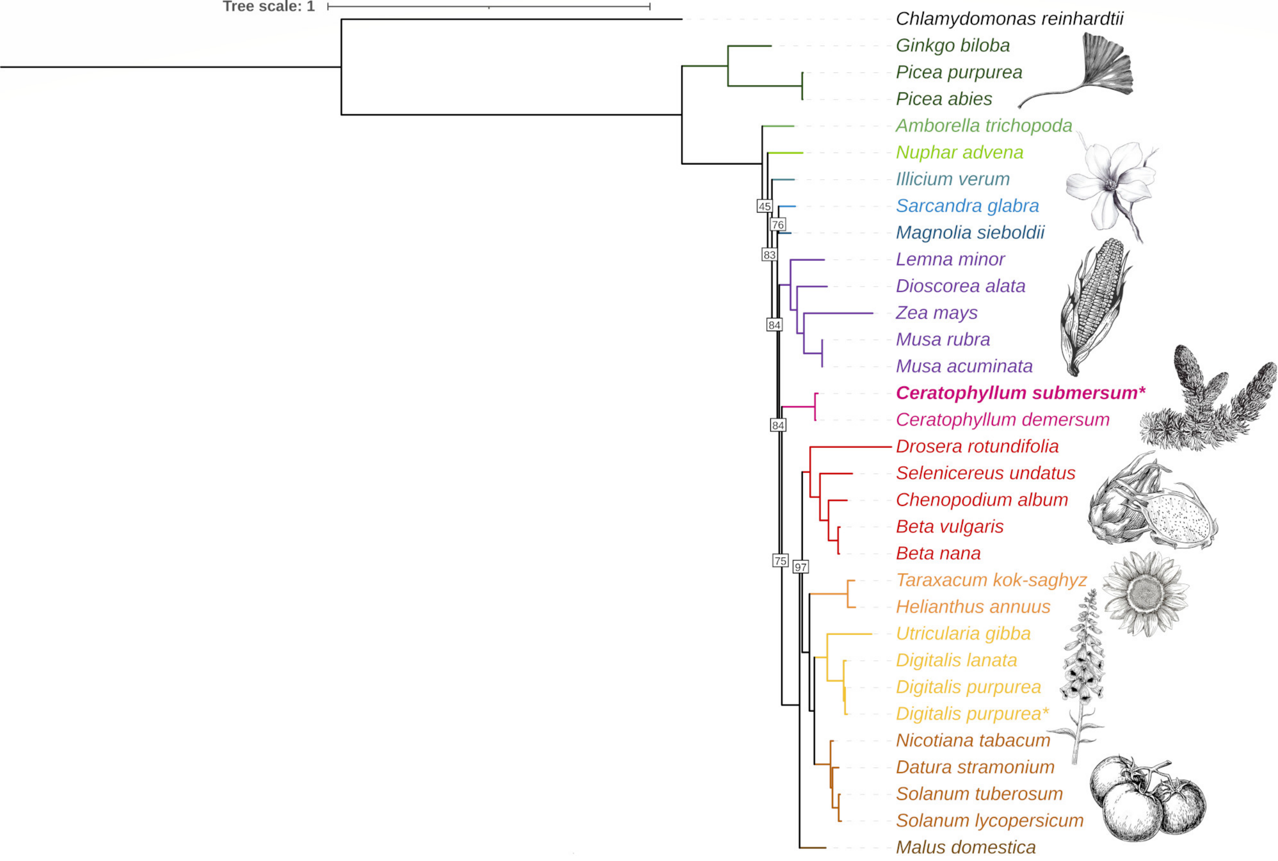

### Additional file 6

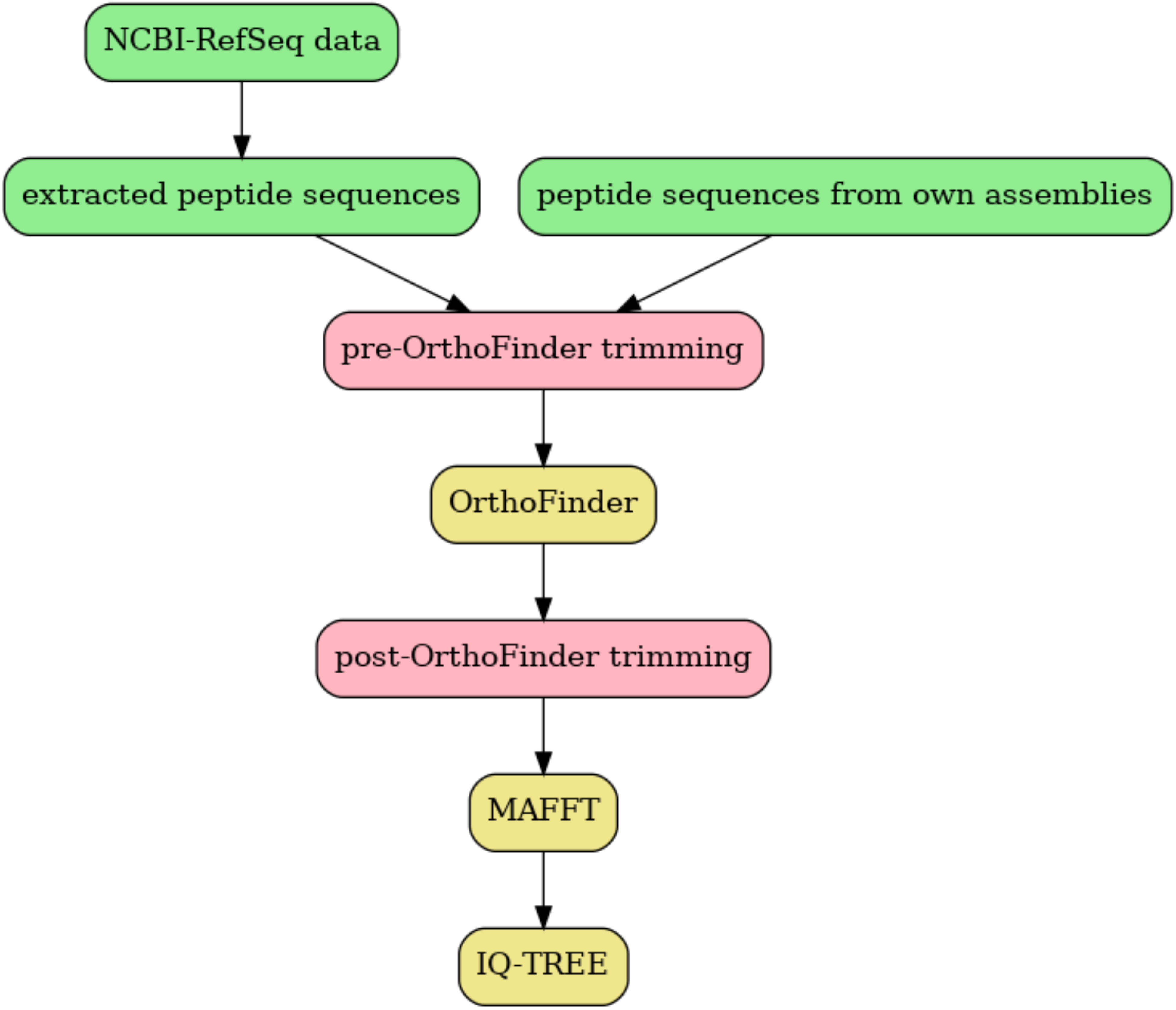
