## Additional file 3 for "Phylogenetic placement of *Ceratophyllum submersum* based on a complete plastome sequence derived from nanopore long read sequencing data"

List of all Reference plastomes from RefSeq used in this study, and their corresponding references.

# NC_001879.2	Nicotiana tabacum
REFERENCE   1  (bases 1 to 155848)
  AUTHORS   Kunnimalaiyaan,M. and Nielsen,B.L.
  TITLE     Fine mapping of replication origins (ori A and ori B) in Nicotiana
            tabacum chloroplast DNA
  JOURNAL   Nucleic Acids Res. 25 (18), 3681-3686 (1997)
   PUBMED   9278490
REFERENCE   2  (bases 1 to 155943)
  AUTHORS   Nadot,S., Bittar,G., Carter,L., Lacroix,R. and Lejeune,B.
  TITLE     A phylogenetic analysis of monocotyledons based on the chloroplast
            gene rps4, using parsimony and a new numerical phenetics method
  JOURNAL   Mol. Phylogenet. Evol. 4 (3), 257-282 (1995)
   PUBMED   8845963
REFERENCE   3  (bases 1 to 155848)
  AUTHORS   Morton,B.R. and Clegg,M.T.
  TITLE     A chloroplast DNA mutational hotspot and gene conversion in a
            noncoding region near rbcL in the grass family (Poaceae)
  JOURNAL   Curr. Genet. 24 (4), 357-365 (1993)
   PUBMED   8252646
REFERENCE   4  (bases 31646 to 32296; 90230 to 91610)
  AUTHORS   Ayliffe,M.A. and Timmis,J.N.
  TITLE     Plastid DNA sequence homologies in the tobacco nuclear genome
  JOURNAL   Mol. Gen. Genet. 236 (1), 105-112 (1992)
   PUBMED   1337369
REFERENCE   5  (bases 1 to 155943)
  AUTHORS   Mitchelson,K. and Stephen,J.
  TITLE     Processing of histidine transfer RNA precursors in tobacco
            chloroplasts
  JOURNAL   Nucleic Acids Res. 19 (11), 3150 (1991)
   PUBMED   2057370
REFERENCE   6  (bases 1 to 155943)
  AUTHORS   Shimada,H. and Sugiura,M.
  TITLE     Fine structural features of the chloroplast genome: comparison of
            the sequenced chloroplast genomes
  JOURNAL   Nucleic Acids Res. 19 (5), 983-995 (1991)
   PUBMED   1708498
REFERENCE   7  (bases 1 to 155848)
  AUTHORS   Shinozaki,K., Ohme,M., Tanaka,M., Wakasugi,T., Hayashida,N.,
            Matsubayashi,T., Zaita,N., Chunwongse,J., Obokata,J.,
            Yamaguchi-Shinozaki,K., Ohto,C., Torazawa,K., Meng,B.Y., Sugita,M.,
            Deno,H., Kamogashira,T., Yamada,K., Kusuda,J., Takaiwa,F., Kato,A.,
            Tohdoh,N., Shimada,H. and Sugiura,M.
  TITLE     The complete nucleotide sequence of the tobacco chloroplast genome:
            its gene organization and expression
  JOURNAL   EMBO J. 5 (9), 2043-2049 (1986)
   PUBMED   16453699
REFERENCE   8  (bases 1 to 155943)
  CONSRTM   NCBI Genome Project
  TITLE     Direct Submission
  JOURNAL   Submitted (30-NOV-2000) National Center for Biotechnology
            Information, NIH, Bethesda, MD 20894, USA
REFERENCE   9  (bases 1 to 155943)
  AUTHORS   Tsudsuki,T.
  TITLE     Direct Submission
  JOURNAL   Submitted (27-FEB-1998) Data Processing Center, Aichi-Gakuin
            University, 12 Araike,1 Iwasaki, Nisshin, Aichi 470-0195, Japan


# NC_039621.1	Sarcandra glabra
REFERENCE   1  (bases 1 to 158900)
  AUTHORS   Zeng,C.X., Hollingsworth,P.M., Yang,J., He,Z.S., Zhang,Z.R.,
            Li,D.Z. and Yang,J.B.
  TITLE     Genome skimming herbarium specimens for DNA barcoding and
            phylogenomics
  JOURNAL   Plant Methods 14, 43 (2018)
   PUBMED   29928291
  REMARK    Publication Status: Online-Only
REFERENCE   2  (bases 1 to 158900)
  CONSRTM   NCBI Genome Project
  TITLE     Direct Submission
  JOURNAL   Submitted (06-NOV-2018) National Center for Biotechnology
            Information, NIH, Bethesda, MD 20894, USA
REFERENCE   3  (bases 1 to 158900)
  AUTHORS   Hollingsworth,P.M., Yang,J., He,Z.-S., Zhang,Z.-R., Li,D.-Z. and
            Yang,J.-B.
  TITLE     Direct Submission
  JOURNAL   Submitted (14-MAY-2018) Germplasm Bank of Wild Species, Kunming
            Institute of Botany, Chinese Academy of Sciences, 132 Lanhei Road,
            Heilongtan, Kunming, Yunnan 65020, China


# NC_001666.2	Zea mays
REFERENCE   1  (bases 1 to 140384)
  AUTHORS   Maier,R.M., Neckermann,K., Igloi,G.L. and Kossel,H.
  TITLE     Complete sequence of the maize chloroplast genome: gene content,
            hotspots of divergence and fine tuning of genetic information by
            transcript editing
  JOURNAL   J. Mol. Biol. 251 (5), 614-628 (1995)
   PUBMED   7666415
REFERENCE   2  (bases 1 to 140384)
  AUTHORS   Weglohner,W. and Subramanian,A.R.
  TITLE     Nucleotide sequence of maize chloroplast rpl32: completing the
            apparent set of plastid ribosomal protein genes and their tentative
            operon organization
  JOURNAL   Plant Mol. Biol. 21 (3), 543-548 (1993)
   PUBMED   8443346
REFERENCE   3  (bases 1 to 140384)
  AUTHORS   Maier,R.M., Neckermann,K., Hoch,B., Akhmedov,N.B. and Kossel,H.
  TITLE     Identification of editing positions in the ndhB transcript from
            maize chloroplasts reveals sequence similarities between editing
            sites of chloroplasts and plant mitochondria
  JOURNAL   Nucleic Acids Res. 20 (23), 6189-6194 (1992)
   PUBMED   1282235
REFERENCE   4  (bases 1 to 140384)
  AUTHORS   Rodermel,S.R.
  TITLE     Nucleotide sequence of a maize chloroplast DNA fragment containing
            an inversion breakpoint, trnG (GCC), trnG (UCC), trnfM, and a trnG
            pseudogene
  JOURNAL   Nucleic Acids Res. 20 (21), 5844 (1992)
   PUBMED   1454547
REFERENCE   5  (bases 1 to 140384)
  AUTHORS   Weglohner,W. and Subramanian,A.R.
  TITLE     Nucleotide sequence of a region of maize chloroplast DNA containing
            the 3' end of clpP, exon 1 of rps12 and rpl20 and their
            cotranscription
  JOURNAL   Plant Mol. Biol. 18 (2), 415-418 (1992)
   PUBMED   1732000
REFERENCE   6  (bases 1 to 140384)
  AUTHORS   Kangasjarvi,J., McCullough,A. and Gengenbach,B.G.
  TITLE     Nucleotide sequence and transcription of maize plastid genome Bam
            HI fragment 14 containing ORF170
  JOURNAL   Plant Mol. Biol. 17 (3), 513-515 (1991)
   PUBMED   1884003
REFERENCE   7  (bases 1 to 140384)
  AUTHORS   Kavousi,M., Giese,K., Larrinua,I.M., McLaughlin,W.E. and
            Subramanian,A.R.
  TITLE     Nucleotide sequence and map positions of the duplicated gene for
            maize (Zea mays) chloroplast ribosomal protein L2
  JOURNAL   Nucleic Acids Res. 18 (14), 4244 (1990)
   PUBMED   2377464
REFERENCE   8  (bases 1 to 140384)
  AUTHORS   Fitzky,B. and Subramanian,A.R.
  TITLE     Nucleotide sequence and map positions of the duplicated gene for
            chloroplast ribosomal protein S15 in Zea mays (maize)
  JOURNAL   Nucleic Acids Res. 18 (11), 3407 (1990)
   PUBMED   2129550
REFERENCE   9  (bases 1 to 140384)
  AUTHORS   Igloi,G.L., Meinke,A., Dory,I. and Kossel,H.
  TITLE     Nucleotide sequence of the maize chloroplast rpo B/C1/C2 operon:
            comparison between the derived protein primary structures from
            various organisms with respect to functional domains
  JOURNAL   Mol. Gen. Genet. 221 (3), 379-394 (1990)
   PUBMED   2381419
REFERENCE   10 (bases 1 to 140384)
  AUTHORS   Steinmuller,K., Ley,A.C., Steinmetz,A.A., Sayre,R.T. and Bogorad,L.
  TITLE     Characterization of the ndhC-psbG-ORF157/159 operon of maize
            plastid DNA and of the cyanobacterium Synechocystis sp. PCC6803
  JOURNAL   Mol. Gen. Genet. 216 (1), 60-69 (1989)
   PUBMED   2499764
REFERENCE   11 (bases 1 to 140384)
  AUTHORS   Bowman,C.M., Barker,R.F. and Dyer,T.A.
  TITLE     In wheat ctDNA, segments of ribosomal protein genes are dispersed
            repeats, probably conserved by nonreciprocal recombination
  JOURNAL   Curr. Genet. 14 (2), 127-136 (1988)
   PUBMED   3180271
REFERENCE   12 (bases 1 to 140384)
  AUTHORS   Ruf,M. and Kossel,H.
  TITLE     Structure and expression of the gene coding for the alpha-subunit
            of DNA-dependent RNA polymerase from the chloroplast genome of Zea
            mays
  JOURNAL   Nucleic Acids Res. 16 (13), 5741-5754 (1988)
   PUBMED   3399379
REFERENCE   13 (bases 1 to 140384)
  AUTHORS   Lukens,J.H. and Bogorad,L.
  TITLE     Nucleotide sequence containing the maize chloroplast proline (UGG)
            and tryptophan (CCA) tRNA genes
  JOURNAL   Nucleic Acids Res. 16 (11), 5192 (1988)
   PUBMED   3387224
REFERENCE   14 (bases 1 to 140384)
  AUTHORS   Schantz,R. and Bogorad,L.
  TITLE     Maize chloroplast genes ndhD, ndhE, and psaC. Sequences,
            transcripts and transcript pools
  JOURNAL   Plant Mol. Biol. 11 (3), 239-247 (1988)
   PUBMED   24272337
REFERENCE   15 (bases 1 to 140384)
  AUTHORS   Giese,K., Subramanian,A.R., Larrinua,I.M. and Bogorad,L.
  TITLE     Nucleotide sequence, promoter analysis, and linkage mapping of the
            unusually organized operon encoding ribosomal proteins S7 and S12
            in maize chloroplast
  JOURNAL   J. Biol. Chem. 262 (31), 15251-15255 (1987)
   PUBMED   2822717
REFERENCE   16 (bases 1 to 140384)
  AUTHORS   Rodermel,S., Orlin,P. and Bogorad,L.
  TITLE     The transcription termination region between two
            convergently-transcribed photoregulated operons in the maize
            plastid chromosome contains rps14, trnR (UCU) and a putative trnfM
            pseudogene
  JOURNAL   Nucleic Acids Res. 15 (13), 5493 (1987)
   PUBMED   3601681
REFERENCE   17 (bases 1 to 140384)
  AUTHORS   McLaughlin,W.E. and Larrinua,I.M.
  TITLE     The sequence of the maize plastid encoded rpl 22 locus
  JOURNAL   Nucleic Acids Res. 15 (10), 4356 (1987)
   PUBMED   3295778
REFERENCE   18 (bases 1 to 140384)
  AUTHORS   Rock,C.D., Barkan,A. and Taylor,W.C.
  TITLE     The maize plastid psbB-psbF-petB-petD gene cluster: spliced and
            unspliced petB and petD RNAs encode alternative products
  JOURNAL   Curr. Genet. 12 (1), 69-77 (1987)
   PUBMED   2835175
REFERENCE   19 (bases 1 to 140384)
  AUTHORS   Erion,J.L.
  TITLE     Characterization of the mRNA transcripts of the maize,
            ribulose-1,5-bisphosphate carboxylase, large subunit gene
  JOURNAL   Plant Mol. Biol. 4 (2-3), 169-179 (1985)
   PUBMED   24310754
REFERENCE   20 (bases 1 to 140384)
  AUTHORS   Fish,L.E., Kuck,U. and Bogorad,L.
  TITLE     Two partially homologous adjacent light-inducible maize chloroplast
            genes encoding polypeptides of the P700 chlorophyll a-protein
            complex of photosystem I
  JOURNAL   J. Biol. Chem. 260 (3), 1413-1421 (1985)
   PUBMED   3881431
REFERENCE   21 (bases 1 to 140384)
  AUTHORS   Strittmatter,G. and Kossel,H.
  TITLE     Cotranscription and processing of 23S, 4.5S and 5S rRNA in
            chloroplasts from Zea mays
  JOURNAL   Nucleic Acids Res. 12 (20), 7633-7647 (1984)
   PUBMED   6093045
REFERENCE   22 (bases 1 to 140384)
  CONSRTM   NCBI Genome Project
  TITLE     Direct Submission
  JOURNAL   Submitted (30-NOV-2000) National Center for Biotechnology
            Information, NIH, Bethesda, MD 20894, USA
REFERENCE   23 (bases 1 to 140384)
  AUTHORS   Neckermann,K.
  TITLE     Direct Submission
  JOURNAL   Submitted (06-SEP-2000) Institut f. Biologie III, Schaenzlestr. 1
            D- 79104, Germany
REFERENCE   24 (bases 1 to 140384)
  AUTHORS   Neckermann,K.
  TITLE     Direct Submission
  JOURNAL   Submitted (25-APR-1995) K. Neckermann, Institut f. Biologie III,
            Schaenzlestr. 1, D- 79104 Freiburg i.Br., FRG
  REMARK    revised by [22]


# NC_021456.1	Picea abies
REFERENCE   1  (bases 1 to 124084)
  AUTHORS   Nystedt,B., Street,N.R., Wetterbom,A., Zuccolo,A., Lin,Y.C.,
            Scofield,D.G., Vezzi,F., Delhomme,N., Giacomello,S., Alexeyenko,A.,
            Vicedomini,R., Sahlin,K., Sherwood,E., Elfstand,M., Gramzow,L.,
            Holmberg,K., Hallman,J., Keech,O., Klasson,L., Koriabine,M.,
            Kucukoglu,M., Kaller,M., Lutman,J., Lysholm,F., Niittyla,T.,
            Olson,A., Rilakovic,N., Ritland,C., Rosello,J.A., Sena,J.,
            Svensson,T., Talavera-Lopez,C., Theissen,G., Tuominen,H.,
            Vanneste,K., Wu,Z.Q., Zhang,B., Zerbe,P., Arvestad,L., Bhalero,R.,
            Bohlmann,J., Bousquet,J., Garcia Gil,R., Hvidsten,T.R., de Jong,P.,
            MacKay,J., Morgante,M., Ritland,K., Sundberg,B., Lee Thompson,S.,
            Van de Peer,Y., Andersson,B., Nilsson,O., Ingvarsson,P.,
            Lundeberg,J. and Jansson,S.
  TITLE     The Norway spruce genome sequence and conifer genome evolution
  JOURNAL   Nature 497 (7451), 579-584 (2013)
   PUBMED   23698360
REFERENCE   2  (bases 1 to 124084)
  CONSRTM   NCBI Genome Project
  TITLE     Direct Submission
  JOURNAL   Submitted (07-JUN-2013) National Center for Biotechnology
            Information, NIH, Bethesda, MD 20894, USA
REFERENCE   3  (bases 1 to 124084)
  AUTHORS   Nystedt,B.
  TITLE     Direct Submission
  JOURNAL   Submitted (12-APR-2013) DBB, Stockholm University, Tomtebodav 23A,
            17165, SWEDEN


# NC_010109.1	Lemna minor
REFERENCE   1  (bases 1 to 165955)
  AUTHORS   Mardanov,A.V., Ravin,N.V., Kuznetsov,B.B., Samigullin,T.H.,
            Antonov,A.S., Kolganova,T.V. and Skyabin,K.G.
  TITLE     Complete sequence of the duckweed (Lemna minor) chloroplast genome:
            structural organization and phylogenetic relationships to other
            angiosperms
  JOURNAL   J. Mol. Evol. 66 (6), 555-564 (2008)
   PUBMED   18463914
REFERENCE   2  (bases 1 to 165955)
  CONSRTM   NCBI Genome Project
  TITLE     Direct Submission
  JOURNAL   Submitted (05-DEC-2007) National Center for Biotechnology
            Information, NIH, Bethesda, MD 20894, USA
REFERENCE   3  (bases 1 to 165955)
  AUTHORS   Ravin,N.V.
  TITLE     Direct Submission
  JOURNAL   Submitted (13-FEB-2006) Center 'Bioengineering' of The Russian
            Academy of Sciences, Prosp 60-let Oktiabria, bld. 7-1, Moscow
            117312, Russia


# NC_018117.1	Datura stramonium
REFERENCE   1  (bases 1 to 155871)
  AUTHORS   Li,X.W., Li,Q., Chen,X.C., Lin,X.H. and Chen,S.L.
  TITLE     High-throughput multiplex sequencing reveals the feasibility of
            chloroplast genomes as a plant super-barcode
  JOURNAL   Unpublished
REFERENCE   2  (bases 1 to 155871)
  CONSRTM   NCBI Genome Project
  TITLE     Direct Submission
  JOURNAL   Submitted (10-JUL-2012) National Center for Biotechnology
            Information, NIH, Bethesda, MD 20894, USA
REFERENCE   3  (bases 1 to 155871)
  AUTHORS   Li,X.W., Li,Q., Chen,X.C., Lin,X.H. and Chen,S.L.
  TITLE     Direct Submission
  JOURNAL   Submitted (07-SEP-2011) Institute of Medicinal Plant Development,
            Chinese Academy of Medical Sciences, No. 151 Malianwa North Road,
            Haidian District, Beijing 100193, P. R. China


# NC_009962.1	Ceratophyllum demersum
REFERENCE   1  (bases 1 to 156252)
  AUTHORS   Moore,M.J., Bell,C.D., Soltis,P.S. and Soltis,D.E.
  TITLE     Using plastid genome-scale data to resolve enigmatic relationships
            among basal angiosperms
  JOURNAL   Proc. Natl. Acad. Sci. U.S.A. 104 (49), 19363-19368 (2007)
   PUBMED   18048334
REFERENCE   2  (bases 1 to 156252)
  CONSRTM   NCBI Genome Project
  TITLE     Direct Submission
  JOURNAL   Submitted (07-NOV-2007) National Center for Biotechnology
            Information, NIH, Bethesda, MD 20894, USA
REFERENCE   3  (bases 1 to 156252)
  AUTHORS   Moore,M.J., Bell,C.D., Soltis,P.S. and Soltis,D.E.
  TITLE     Direct Submission
  JOURNAL   Submitted (16-MAY-2007) Department of Botany, University of
            Florida, PO Box 117800, Gainesville, FL 32611-7800, USA


# NC_034950.1	Chenopodium album
REFERENCE   1  (bases 1 to 152167)
  AUTHORS   Cho,K.-S., Cheon,K.-S., Yoo,K.-O., Hong,S.-Y. and Lee,H.-O.
  TITLE     Comparative analysis of Chenopodium quinoa and C. album by complete
            chloroplast genome sequences
  JOURNAL   Unpublished
REFERENCE   2  (bases 1 to 152167)
  CONSRTM   NCBI Genome Project
  TITLE     Direct Submission
  JOURNAL   Submitted (02-JUN-2017) National Center for Biotechnology
            Information, NIH, Bethesda, MD 20894, USA
REFERENCE   3  (bases 1 to 152167)
  AUTHORS   Cho,K.-S., Cheon,K.-S., Yoo,K.-O., Hong,S.-Y. and Lee,H.-O.
  TITLE     Direct Submission
  JOURNAL   Submitted (03-JAN-2017) Highland Agriculture Research Institute,
            Rural Development Administration, Honggye-Ri, Pyeongchang-Gun,
            Gangwon-Do 232-955, Republic of Korea


# NC_021449.1	Utricularia gibba
REFERENCE   1  (bases 1 to 152113)
  AUTHORS   Ibarra-Laclette,E., Lyons,E., Hernandez-Guzman,G.,
            Perez-Torres,C.A., Carretero-Paulet,L., Chang,T.H., Lan,T.,
            Welch,A.J., Juarez,M.J., Simpson,J., Fernandez-Cortes,A.,
            Arteaga-Vazquez,M., Gongora-Castillo,E., Acevedo-Hernandez,G.,
            Schuster,S.C., Himmelbauer,H., Minoche,A.E., Xu,S., Lynch,M.,
            Oropeza-Aburto,A., Cervantes-Perez,S.A., de Jesus
            Ortega-Estrada,M., Cervantes-Luevano,J.I., Michael,T.P.,
            Mockler,T., Bryant,D., Herrera-Estrella,A., Albert,V.A. and
            Herrera-Estrella,L.
  TITLE     Architecture and evolution of a minute plant genome
  JOURNAL   Nature 498 (7452), 94-98 (2013)
   PUBMED   23665961
REFERENCE   2  (bases 1 to 152113)
  CONSRTM   NCBI Genome Project
  TITLE     Direct Submission
  JOURNAL   Submitted (07-JUN-2013) National Center for Biotechnology
            Information, NIH, Bethesda, MD 20894, USA
REFERENCE   3  (bases 1 to 152113)
  AUTHORS   Ibarra-Laclette,E., Hernandez-Guzman,G., Perez-Torres,C.A.,
            Carretero-Paulet,L., Chang,T.-H., Lan,T., Welch,A.J., Abraham
            Juarez,M.J., Simpson,J., Fernandez-Cortes,A., Arteaga-Vazquez,M.,
            Gongora-Castillo,E., Acevedo-Hernandez,G., Schuster,S.C.,
            Himmelbauer,H., Minoche,A.E., Xu,S., Lynch,M., Oropeza-Aburto,A.,
            Cervantes-Perez,S.A., de Jesus Ortega-Estrada,M.,
            Cervantes-Luevano,J.I., Michael,T.P., Mockler,T., Bryant,D.,
            Herrera-Estrella,A., Albert,V.A. and Herrera-Estrella,L.
  TITLE     Direct Submission
  JOURNAL   Submitted (11-APR-2013) Department of Biological Sciences,
            University at Buffalo, 109 Cooke Hall, Buffalo, NY 14260, USA


# NC_029770.1	Drosera rotundifolia
REFERENCE   1  (bases 1 to 192912)
  AUTHORS   Mardanov,A.V., Beletsky,A.V., Kadnikov,V.V. and Ravin,N.V.
  TITLE     Complete chloroplast genome of Droesria rothundifolia
  JOURNAL   Unpublished
REFERENCE   2  (bases 1 to 192912)
  CONSRTM   NCBI Genome Project
  TITLE     Direct Submission
  JOURNAL   Submitted (24-MAR-2016) National Center for Biotechnology
            Information, NIH, Bethesda, MD 20894, USA
REFERENCE   3  (bases 1 to 192912)
  AUTHORS   Mardanov,A.V., Gruzdev,E.V., Kadnikov,V.V., Beletsky,A.V. and
            Ravin,N.N.
  TITLE     Direct Submission
  JOURNAL   Submitted (17-NOV-2015) Institute of Bioengineering, Research
            Center of Biotechnology of RAS, Prosp. 60-let Oktyabrya, bld. 7-1,
            Moscow 117312, Russia


# NC_068046.1	Digitalis purpurea
REFERENCE   1  (bases 1 to 152999)
  AUTHORS   Zhao,F., Liu,B., Liu,S., Min,D.-Z., Zhang,T., Cai,J., Zhou,X.-X.,
            Chen,B., Olmstead,R.G., Xiang,C.-L. and Li,B.
  TITLE     Disentangling a 40-year-old taxonomic puzzle: the phylogenetic
            position of Mimulicalyx (Lamiales)
  JOURNAL   Bot J Linn Soc (2022)
  REMARK    DOI: 10.1093/botlinnean/boac033
REFERENCE   2  (bases 1 to 152999)
  CONSRTM   NCBI Genome Project
  TITLE     Direct Submission
  JOURNAL   Submitted (12-NOV-2022) National Center for Biotechnology
            Information, NIH, Bethesda, MD 20894, USA
REFERENCE   3  (bases 1 to 152999)
  AUTHORS   Zhao,F., Liu,B., Liu,S., Min,D.-Z., Zhang,T., Cai,J., Zhou,X.-X.,
            Chen,B., Olmstead,R.G., Xiang,C.-L. and Li,B.
  TITLE     Direct Submission
  JOURNAL   Submitted (17-DEC-2021) Key Laboratory for Plant Diversity and
            Biogeography of East Asia, Kunming Institute of Botany, Chinese
            Academy of Sciences, Panlong District, Lanhei Road 132, Kunming
            City, Yunnan 650201, China


# NC_039707.1	Dioscorea alata
REFERENCE   1  (bases 1 to 153161)
  AUTHORS   Cao,J., Jiang,D., Zhao,Z., Yuan,S., Zhang,Y., Zhang,T., Zhong,W.,
            Yuan,Q. and Huang,L.
  TITLE     Development of Chloroplast Genomic Resources in Chinese Yam
            (Dioscorea polystachya)
  JOURNAL   Biomed Res Int 2018, 1-11 (2018)
REFERENCE   2  (bases 1 to 153161)
  CONSRTM   NCBI Genome Project
  TITLE     Direct Submission
  JOURNAL   Submitted (09-NOV-2018) National Center for Biotechnology
            Information, NIH, Bethesda, MD 20894, USA
REFERENCE   3  (bases 1 to 153161)
  AUTHORS   Yuan,Q.-J.
  TITLE     Direct Submission
  JOURNAL   Submitted (25-OCT-2017) Center for Chinese Materia Medica
            Resources, China Academy of Chinese Medical Sciences, No 16 Nan
            xiao jie, Dong zhi men nei, Beijing, Beijing 100700, China


# NC_064363.1	Picea purpurea
REFERENCE   1  (bases 1 to 124233)
  CONSRTM   NCBI Genome Project
  TITLE     Direct Submission
  JOURNAL   Submitted (23-JUN-2022) National Center for Biotechnology
            Information, NIH, Bethesda, MD 20894, USA
REFERENCE   2  (bases 1 to 124233)
  AUTHORS   Wang,C.
  TITLE     Direct Submission
  JOURNAL   Submitted (16-NOV-2021) Agriculture and Animal Husbandry, Qinghai
            University, Ningda Road, Xining, Qinghai 810016, China


# NC_061549.1	Malus domestica
REFERENCE   1  (bases 1 to 160288)
  AUTHORS   Tian,X.
  TITLE     Complete chloroplast genomes studies of different apple varieties
            indicated modern cultivated apples originate from Malus sieversii
            and Malus sylvestris
  JOURNAL   Unpublished
REFERENCE   2  (bases 1 to 160288)
  CONSRTM   NCBI Genome Project
  TITLE     Direct Submission
  JOURNAL   Submitted (31-MAR-2022) National Center for Biotechnology
            Information, NIH, Bethesda, MD 20894, USA
REFERENCE   3  (bases 1 to 160288)
  AUTHORS   Tian,X.
  TITLE     Direct Submission
  JOURNAL   Submitted (07-OCT-2021) College of Life Science and Technology,
            Xinjiang University, 666 Shengli Road, Urumqi, Xinjiang, China,
            Urumqi, Xinjiang 830046, China


# NC_008788.1	Nuphar advena
REFERENCE   1  (bases 1 to 160866)
  AUTHORS   Raubeson,L.A., Peery,R., Chumley,T.W., Dziubek,C., Fourcade,H.M.,
            Boore,J.L. and Jansen,R.K.
  TITLE     Comparative chloroplast genomics: analyses including new sequences
            from the angiosperms Nuphar advena and Ranunculus macranthus
  JOURNAL   BMC Genomics 8, 174 (2007)
   PUBMED   17573971
  REMARK    Publication Status: Online-Only
REFERENCE   2  (bases 1 to 160866)
  CONSRTM   NCBI Genome Project
  TITLE     Direct Submission
  JOURNAL   Submitted (11-JAN-2007) National Center for Biotechnology
            Information, NIH, Bethesda, MD 20894, USA
REFERENCE   3  (bases 1 to 160866)
  AUTHORS   Raubeson,L., Chumley,T., Peery,R., Dziubek,C., Fourcade,M.,
            Boore,J. and Jansen,R.
  TITLE     Direct Submission
  JOURNAL   Submitted (06-JAN-2006) Biological Sciences, Central Washington
            University, 400 E University Way, Ellensburg, WA 98926-7537, USA


# NC_008096.2	Solanum tuberosum
REFERENCE   1  (bases 1 to 155296)
  AUTHORS   Gargano,D., Vezzi,A., Scotti,N., Gray,J.C., Valle,G., Grillo,S. and
            Cardi,T.
  TITLE     The complete nucleotide sequence of potato (Solanum tuberosum cv.
            Desiree) chloroplast DNA
  JOURNAL   (in) ? (Ed.); ABSTRACTS 2ND SOLANACEAE GENOME WORKSHOP 2005: 107;
            (2005)
REFERENCE   2  (bases 1 to 155296)
  AUTHORS   Gargano,D., Scotti,N., Vezzi,A., Bilardi,A., Valle,G., Grillo,S.
            and Cardi,T.
  TITLE     Complete chloroplast genome sequences of Solanum tuberosum cultivar
            Desiree and comparative analyses with other Solanaceae genomes
  JOURNAL   Unpublished
REFERENCE   3  (bases 1 to 155296)
  CONSRTM   NCBI Genome Project
  TITLE     Direct Submission
  JOURNAL   Submitted (09-JUN-2006) National Center for Biotechnology
            Information, NIH, Bethesda, MD 20894, USA
REFERENCE   4  (bases 1 to 155296)
  AUTHORS   Gargano,D., Scotti,N., Vezzi,A., Bilardi,A., Valle,G., Grillo,S.
            and Cardi,T.
  TITLE     Direct Submission
  JOURNAL   Submitted (01-FEB-2006) CNR-IGV, Institute of Plant Genetics, Via
            Universita' 133, Portici 80055, Italy


# NC_032057.1	Taraxacum kok-saghyz
REFERENCE   1  (bases 1 to 151338)
  AUTHORS   Zhang,Y., Iaffaldano,B.J., Zhuang,X., Cardina,J. and Cornish,K.
  TITLE     Chloroplast genome resources and molecular markers differentiate
            rubber dandelion species from weedy relatives
  JOURNAL   Unpublished
REFERENCE   2  (bases 1 to 151338)
  CONSRTM   NCBI Genome Project
  TITLE     Direct Submission
  JOURNAL   Submitted (07-DEC-2016) National Center for Biotechnology
            Information, NIH, Bethesda, MD 20894, USA
REFERENCE   3  (bases 1 to 151338)
  AUTHORS   Zhang,Y., Iaffaldano,B.J., Zhuang,X., Cardina,J. and Cornish,K.
  TITLE     Direct Submission
  JOURNAL   Submitted (05-MAY-2016) Department of Horticulture and Crop
            Science, The Ohio State University, Ohio Agricultural Research and
            Development Center, 1680 Madison Avenue, Wooster, OH 44691, USA


# NC_005353.1	Chlamydomonas reinhardtii
REFERENCE   1  (bases 1 to 203828)
  AUTHORS   Maul,J.E., Lilly,J.W., Cui,L., dePamphilis,C.W., Miller,W.,
            Harris,E.H. and Stern,D.B.
  TITLE     The Chlamydomonas reinhardtii plastid chromosome: islands of genes
            in a sea of repeats
  JOURNAL   Plant Cell 14 (11), 2659-2679 (2002)
   PUBMED   12417694
REFERENCE   2  (bases 1 to 203828)
  CONSRTM   NCBI Genome Project
  TITLE     Direct Submission
  JOURNAL   Submitted (21-SEP-2004) National Center for Biotechnology
            Information, NIH, Bethesda, MD 20894, USA
REFERENCE   3  (bases 1 to 203828)
  AUTHORS   Maul,J.E., Lilly,J.W. and Stern,D.B.
  TITLE     Direct Submission
  JOURNAL   Submitted (10-JUL-2003) Plant Molecular Biology, Boyce Thompson
            Institute for Plant Research, Tower Rd., Ithaca, NY 14853, USA
  REMARK    Sequence update by submitter
REFERENCE   4  (bases 1 to 203828)
  AUTHORS   Maul,J.E., Lilly,J.W. and Stern,D.B.
  TITLE     Direct Submission
  JOURNAL   Submitted (20-AUG-2002) Plant Molecular Biology, Boyce Thompson
            Institute for Plant Research, Tower Rd., Ithaca, NY 14853, USA


# NC_065468.1	Musa acuminata AAA Group
REFERENCE   1  (bases 1 to 170299)
  AUTHORS   Shekhawat,U., Visendi,P., Paul,J.-Y., Jung,H., Hamill,S.,
            Tushemereirwe,W., Harding,R. and Dale,J.
  TITLE     Chloroplast genome of Musa acuminata cv. Grand Nain
  JOURNAL   Unpublished
REFERENCE   2  (bases 1 to 170299)
  CONSRTM   NCBI Genome Project
  TITLE     Direct Submission
  JOURNAL   Submitted (10-AUG-2022) National Center for Biotechnology
            Information, NIH, Bethesda, MD 20894, USA
REFERENCE   3  (bases 1 to 170299)
  AUTHORS   Shekhawat,U., Visendi,P., Paul,J.-Y., Jung,H., Hamill,S.,
            Tushemereirwe,W., Harding,R. and Dale,J.
  TITLE     Direct Submission
  JOURNAL   Submitted (23-JUN-2021) Centre for Agriculture and the Bioeconomy,
            Queensland University of Technology, 2 George Street, Brisbane, Qld
            4000, Australia


# NC_007898.3	Solanum lycopersicum
REFERENCE   1  (bases 1 to 155461)
  AUTHORS   Daniell,H., Lee,S.-B., Grevich,J., Saski,C., Quesada-Vargas,T.,
            Guda,C., Tomkins,J. and Jansen,R.K.
  TITLE     Complete chloroplast genome sequences of Solanum tuberosum, Solanum
            lycopersicum and comparative analyses with other Solanaceae genomes
  JOURNAL   Unpublished
REFERENCE   2  (bases 1 to 155461)
  CONSRTM   NCBI Genome Project
  TITLE     Direct Submission
  JOURNAL   Submitted (17-SEP-2013) National Center for Biotechnology
            Information, NIH, Bethesda, MD 20894, USA
REFERENCE   3  (bases 1 to 155461)
  AUTHORS   Daniell,H., Lee,S.-B., Grevich,J., Saski,C., Quesada-Vargas,T.,
            Guda,C., Tomkins,J. and Jansen,R.K.
  TITLE     Direct Submission
  JOURNAL   Submitted (03-JAN-2006) Molecular Biology & Microbiology,
            University of Central Florida, Biomolecular Science, Building #20,
            Orlando, FL 32816-2364, USA


# NC_007977.1	Helianthus annuus
REFERENCE   1  (bases 1 to 151104)
  AUTHORS   Timme,R.E., Kuehl,J.V., Boore,J.L. and Jansen,R.K.
  TITLE     A comparison of the first two published chloroplast genomes in
            Asteraceae: Lactuca and Helianthus
  JOURNAL   Unpublished
REFERENCE   2  (bases 1 to 151104)
  CONSRTM   NCBI Genome Project
  TITLE     Direct Submission
  JOURNAL   Submitted (03-MAY-2006) National Center for Biotechnology
            Information, NIH, Bethesda, MD 20894, USA
REFERENCE   3  (bases 1 to 151104)
  AUTHORS   Timme,R.E., Kuehl,J.V., Boore,J.L. and Jansen,R.K.
  TITLE     Direct Submission
  JOURNAL   Submitted (30-JAN-2006) Integrative Biology, The University of
            Texas at Austin, 1 University Station, A6700, Austin, TX 78712, USA


# NC_005086.1	Amborella trichopoda
REFERENCE   1
  AUTHORS   Goremykin,V.V., Hirsch-Ernst,K.I., Wolfl,S. and Hellwig,F.H.
  TITLE     Analysis of the Amborella trichopoda Chloroplast Genome Sequence
            Suggests That Amborella Is Not a Basal Angiosperm
  JOURNAL   Mol. Biol. Evol. 20 (9), 1499-1505 (2003)
   PUBMED   12832641
REFERENCE   2  (bases 1 to 162686)
  CONSRTM   NCBI Genome Project
  TITLE     Direct Submission
  JOURNAL   Submitted (08-SEP-2003) National Center for Biotechnology
            Information, NIH, Bethesda, MD 20894, USA
REFERENCE   3  (bases 1 to 162686)
  AUTHORS   Goremykin,V. and Hellwig,F.
  TITLE     Direct Submission
  JOURNAL   Submitted (05-NOV-2002) Institut fuer Spezielle Botanik,
            Friedrich-Schiller-Universitaet Jena, Philosophenweg 16, 07743
            Jena, Germany
REFERENCE   4  (bases 1 to 162686)
  AUTHORS   Goremykin,V. and Hellwig,F.
  TITLE     Direct Submission
  JOURNAL   Submitted (05-AUG-2002) Institut fuer Spezielle Botanik,
            Friedrich-Schiller-Universitaet Jena, Philosophenweg 16, 07743
            Jena, Germany
  REMARK    Revised by [3]


# NC_034688.1	Digitalis lanata
REFERENCE   1  (bases 1 to 153108)
  AUTHORS   Zhang,N., Handy,S.M. and Wen,J.
  TITLE     Plastid Genomes of Illicium
  JOURNAL   Unpublished
REFERENCE   2  (bases 1 to 153108)
  CONSRTM   NCBI Genome Project
  TITLE     Direct Submission
  JOURNAL   Submitted (25-MAY-2017) National Center for Biotechnology
            Information, NIH, Bethesda, MD 20894, USA
REFERENCE   3  (bases 1 to 153108)
  AUTHORS   Zhang,N., Ramachandran,P., Ottesen,A.R., Timme,R.E., Handy,S.M. and
            Wen,J.
  TITLE     Direct Submission
  JOURNAL   Submitted (31-OCT-2016) Center of Food Safety and Nutrition, Food
            and Drug Adminstration, 5001 Campus Drive, College Park, MD 20740,
            USA


# NC_016986.1	Ginkgo biloba
REFERENCE   1  (bases 1 to 156988)
  AUTHORS   Li,X., Li,Q., Lin,X., Hu,Z. and Chen,S.
  TITLE     High-throughput multiplex sequencing reveals the prospect of
            chloroplast genomes as a plant super-barcode
  JOURNAL   Unpublished
REFERENCE   2  (bases 1 to 156988)
  CONSRTM   NCBI Genome Project
  TITLE     Direct Submission
  JOURNAL   Submitted (15-MAR-2012) National Center for Biotechnology
            Information, NIH, Bethesda, MD 20894, USA
REFERENCE   3  (bases 1 to 156988)
  AUTHORS   Li,X., Li,Q., Lin,X., Hu,Z. and Chen,S.
  TITLE     Direct Submission
  JOURNAL   Submitted (28-SEP-2011) Resources Center, Institute of Medicinal
            Plant Development, No 151 Malianwa North Road, Beijing, Beijing
            100193, China


# NC_058936.1	Musa rubra
REFERENCE   1  (bases 1 to 169309)
  AUTHORS   Fu,N., Ji,M.-Y. and Ge,X.-G.
  TITLE     Chloroplast genomes
  JOURNAL   Unpublished
REFERENCE   2  (bases 1 to 169309)
  CONSRTM   NCBI Genome Project
  TITLE     Direct Submission
  JOURNAL   Submitted (03-NOV-2021) National Center for Biotechnology
            Information, NIH, Bethesda, MD 20894, USA
REFERENCE   3  (bases 1 to 169309)
  AUTHORS   Fu,N., Ji,M.-Y. and Ge,X.-G.
  TITLE     Direct Submission
  JOURNAL   Submitted (02-SEP-2021) Plant science center, South China Botanical
            Garden, No. 723 Xingke Road, Guangzhou, Guangdong 510650, China


# NC_041435.1	Magnolia sieboldii
REFERENCE   1  (bases 1 to 160177)
  AUTHORS   Gao,J., Song,Y. and Zheng,B.
  TITLE     Complete chloroplast genome sequence of an endangered tree species,
            Magnolia sieboldii (Magnoliaceae)
  JOURNAL   Mitochondrial DNA B Resour 3 (2), 1261-1262 (2018)
REFERENCE   2  (bases 1 to 160177)
  CONSRTM   NCBI Genome Project
  TITLE     Direct Submission
  JOURNAL   Submitted (10-APR-2019) National Center for Biotechnology
            Information, NIH, Bethesda, MD 20894, USA
REFERENCE   3  (bases 1 to 160177)
  AUTHORS   Gao,J. Jr. and Song,Y.T. Sr.
  TITLE     Direct Submission
  JOURNAL   Submitted (27-JUN-2018) Center for Integrative Conservation,
            Xishuangbanna Tropical Botanical Garden, Chinese Academy of
            Sciences, Menglun, Mengla, Yunnan 666303, China


# NC_034689.1	Illicium verum
REFERENCE   1  (bases 1 to 143187)
  AUTHORS   Zhang,N. and Handy,S.M.
  TITLE     Plastid Genomes of Illicium
  JOURNAL   Unpublished
REFERENCE   2  (bases 1 to 143187)
  CONSRTM   NCBI Genome Project
  TITLE     Direct Submission
  JOURNAL   Submitted (25-MAY-2017) National Center for Biotechnology
            Information, NIH, Bethesda, MD 20894, USA
REFERENCE   3  (bases 1 to 143187)
  AUTHORS   Zhang,N., Ramachandran,P., Ottesen,A.R., Timme,R.E., Handy,S.M. and
            Luo,Y.
  TITLE     Direct Submission
  JOURNAL   Submitted (31-OCT-2016) Center of Food Safety and Nutrition, Food
            and Drug Adminstration, 5100 Paint Branch Pkwy, College Park, MD
            20740, USA


# NC_053698.1	Selenicereus undatus
REFERENCE   1  (bases 1 to 133326)
  CONSRTM   NCBI Genome Project
  TITLE     Direct Submission
  JOURNAL   Submitted (25-MAR-2021) National Center for Biotechnology
            Information, NIH, Bethesda, MD 20894, USA
REFERENCE   2  (bases 1 to 133326)
  AUTHORS   Niu,Y.Y. and Liu,J.J.
  TITLE     Direct Submission
  JOURNAL   Submitted (14-AUG-2020) Center of molecular biology, Yunnan
            Institute of Tropical Crop, 99 Xuanwei Avenue, Xishuangbanna,
            Yunnan 666100, China
